# Malaria parasite immune evasion and adaptation to its mosquito host is influenced by the acquisition of multiple blood meals

**DOI:** 10.1101/801480

**Authors:** Hyeogsun Kwon, Rebekah A. Reynolds, Maria L. Simões, George Dimopoulos, Ryan C. Smith

## Abstract

A minimum of two blood meals are required for a mosquito to acquire and transmit malaria, yet *Anopheles* mosquitoes frequently obtain additional blood meals during their adult lifespan. To determine the impact of subsequent blood-feeding on parasite development in *Anopheles gambiae*, we examined rodent and human *Plasmodium* parasite infection with or without an additional non-infected blood meal. We find that an additional blood meal significantly reduces *P. berghei* immature oocyst numbers, yet does not influence mature oocysts that have already begun sporogony. This is in contrast to experiments performed with the human parasite, *P. falciparum*, where an additional blood meal does not affect oocyst numbers. These observations are reproduced when mosquitoes were similarly challenged with an artificial protein meal, suggesting that parasite losses are due to the physical distension of the mosquito midgut. We provide evidence that feeding compromises the integrity of the midgut basal lamina, enabling the recognition and lysis of immature *P. berghei* oocysts by the mosquito complement system. Moreover, we demonstrate that additional feeding promotes *P. falciparum* oocyst growth, suggesting that human malaria parasites exploit host resources provided with blood-feeding to accelerate their growth. This contrasts experiments with *P. berghei*, where the size of surviving oocysts is independent of an additional blood meal. Together, these data demonstrate differences in the ability of *Plasmodium* species to evade immune detection and adapt to utilize host resources at the oocyst stage, representing an additional, yet unexplored component of vectorial capacity that has important implications for transmission of malaria.

## Introduction

Blood-feeding is an inherent behavior of all hematophagous arthropods, providing nutritional resources for development or reproduction, while enabling the acquisition and transmission of a pathogen from one host to the next. This includes a number of arthropod-borne diseases that influence human health, most notably malaria, which causes more than 200 million infections and 400,000 deaths every year (WHO, 2018). Caused by *Plasmodium* parasites, malaria transmission requires the bite of an *Anopheles* mosquito, such that understanding the factors that influence vectorial capacity are integral to efforts to reduce malaria transmission.

Following the ingestion of an infectious blood-meal, malaria parasites undergo substantial development in the mosquito host as they transition from gametes to a fertilized zygote, a motile ookinete, an oocyst, and a sporozoite capable of transmission to a new host (Smith et al., 2014). During this approximate two week period of development (referred to as the extrinsic incubation period, EIP), significant bottlenecks reduce parasite numbers at each of these respective *Plasmodium* stages (Smith et al., 2014). These losses are mediated in part by the mosquito innate immune system that target the *Plasmodium* ookinete or oocyst through distinct immune mechanisms (Gupta et al., 2009; Kwon and Smith, 2019; Kwon et al., 2017; Smith et al., 2015). However, for those parasites that are able to escape immune recognition, the impact of changes to mosquito physiology on the remainder of the parasite life cycle remains unknown. This includes nutritional stress (i.e.-starvation and dehydration) and the potential for multiple blood meals before *Plasmodium* sporozoites reach the salivary glands of the mosquito, which could significantly impact the EIP and the likelihood of transmission (Ohm et al., 2018).

With the ability to complete a gonotrophic cycle approximately every three days, mosquitoes can feed multiple times during their lifespan. Therefore, the consequences of additional feeding behaviors following an initial infection are crucial to our understanding of malaria transmission. In limited studies, *Plasmodium*-infected mosquitoes were more likely to seek an additional blood-meal (Ferguson and Read, 2004), while others have demonstrated that additional feeding increased sporozoite infection of the mosquito salivary glands (Ponnudurai et al., 1989; Rosenberg and Rungsiwongse, 1991). However, the impacts of a blood-meal on developing *Plasmodium* parasites have not been fully explored.

Herein, we examine the influence of an additional blood-meal on *Plasmodium* oocyst survival and development. Performing experiments with both rodent and human malaria parasites, we see distinct differences in parasite survival and growth in response to additional feeding. Our results suggest that *P. falciparum* oocysts have evolved mechanisms to evade immune detection and to capture host resources to facilitate their growth, arguing that an additional blood-meal increases the likelihood of malaria transmission. Therefore, these findings provide novel insight into the host-parasite interactions that determine vectorial capacity and define important new implications for the role of mosquito feeding behavior in the efficacy of malaria transmission.

## Materials and Methods

### Ethics statement

The protocols and procedures used in this study were approved by the Animal Care and Use Committee at Iowa State University (IACUC-18-228) and Johns Hopkins University (M006H300), with additional oversight from the Johns Hopkins School of Public Health Ethics Committee. Commercial anonymous human blood was used for parasite cultures and mosquito feeding experiments, therefore human consent was not required.

### Mosquito rearing

*An. gambiae* mosquitoes of the Keele strain (Hurd et al., 2005; Ranford-Cartwright et al., 2016), as well as the TEP1 mutant and parental control X1 lines (Kwon et al., 2017; Smidler et al., 2013) were reared at 27°C with 80% relative humidity and a 14/10 hour light/dark cycle. At Iowa State University, larvae were fed fish flakes (Tetramin, Tetra), while adult mosquitoes were maintained on 10% sucrose solution and fed on commercial sheep blood for egg production. At the Johns Hopkins Bloomberg School of Public Health, larvae were reared on a diet of fish flakes and cat food, while adult mosquitoes were fed on anesthetized 6- to 8-week-old female Swiss Webster mice for egg production. The Keele colony at Iowa State was derived from the Keele colony at Johns Hopkins and has been independently maintained for ∼4 years.

### *Plasmodium berghei* infection

Female Swiss Webster mice were infected with *P. berghei*-mCherry strain as described previously (Kwon et al., 2017; Smith et al., 2015) and maintained at 19°C with 80% relative humidity and a 14/10 hour light/dark cycle. Malaria parasite infection was examined by dissecting individual mosquito midguts in PBS to perform counts of *Plasmodium* oocyst numbers by fluorescence microscopy (Nikon Eclipse 50i, Nikon) at either eight- or ten-days post-infection.

### *Plasmodium falciparum* infection

Three- to four-day-old female mosquitoes were fed through artificial membrane feeders on an NF54 *P. falciparum* gametocyte culture in human blood as previously (Simões et al., 2017). After removal of the unfed females, *P. falciparum*-infected *An. gambiae* females were maintained at 27°C on a 10% sucrose solution. Midguts were dissected in PBS and stained in 0.2% mercurochrome to determine oocyst numbers at eight- or ten-days post-infection using a light-contrast microscope. Images were captured using an optical microscope.

### Additional feeding challenge following *Plasmodium* infection

Naïve female mosquitoes (3- to 6-day old) were initially challenged with a *P. berghei*-infected mouse or a *P. falciparum*-infected blood meal. Infected mosquitoes were provided with an egg cup for oviposition, then separated into two groups, one of which was maintained on a 10% sucrose solution for the duration of the experiment. The second group was challenged with either defibrinated sheep blood, human blood, or a protein meal (consisting of 200 mg/mL of bovine serum albumin, 2 mM ATP, and 20% (v/v) food dye in 1XPBS) using a glass membrane feeder at either four- or eight-days post-infection to examine the effects of an additional feeding on early or mature oocysts.

Additional experiments performed with the TEP1 mutant and X1 lines were similarly initially infected with *P. berghei*, then challenged with a naïve mouse at day 4 post-infection or maintained on a sucrose diet without receiving a second blood meal. Parasite numbers were evaluated by counting oocyst numbers at day 8 post-infection.

### Measurement of basal lamina integrity using collagen hybridizing peptide (CHP)

Following blood or protein feeding, midguts were dissected from mosquitoes after feeding blood meal or a protein meal at 3, 6, 18, 24, and 48 hrs. Midguts from non-fed mosquitoes served as negative control, while heat-treated midguts (70°C for 10 min) were used as a positive control. After dissection, the blood or protein bolus was removed, washed in 1xPBS, then fixed with 2% glutaraldehyde and 2% paraformaldehyde in PBS at pH 7.4 (Electron Microscopy Sciences) for 3 h at 4°C. Following fixation, samples were washed three times in 1xPBS, then blocked overnight at 4°C in blocking buffer (5% bovine serum albumin in 1xPBS). To measure the potential disruption of collagen present on the midgut basal lamina, fluorescein conjugated collagen hybridizing peptide (CHP) (Echelon Biosciences) was diluted in 1xPBS (1:20; 5 µM). Before use, the CHP dilution was placed on a heating block at 80°C for 10 min, then chilled on ice. CHP was added to the blocked midgut samples and incubated overnight at 4°C. Midguts were washed five times in PBS, then mounted with ProLong®Diamond Antifade mountant with DAPI (Life Technologies). Staining was visualized by fluorescence on a Nikon Eclipse 50i and captured using NIS Elements (Nikon) imaging software under the same exposure settings. Micrographs were used to quantify fluorescence across samples using Image J software (Schneider et al., 2012).

### Immunofluorescence assays

Mosquitoes previously infected with *P. berghei* were either maintained on 10% sucrose or fed on defibrinated sheep blood at day 4 post-infection. Midguts were dissected from ∼24 h after additional blood feeding or from non-challenged mosquitoes. Midguts sheets were prepared, removing the blood bolus. Midgut samples were washed in 1xPBS before fixation in 4% PFA for 1 hour at RT. To examine basal lamina integrity based on staining of the oocyst capsule, midgut samples were washed three times in 1x PBS then blocked overnight in 1% bovine serum albumin (BSA)/ 0.1% Triton X-100 in 1xPBS at 4°C. Midgut samples were incubated with mouse circumsporozoite protein (CSP, 1:500) and rabbit-TEP1 (1:500) primary antibodies overnight in blocking buffer (1% BSA/1xPBS) at 4°C. After washing in 1xPBS, midguts were incubated with Alexa Fluor 488 goat anti-mouse IgG (1:500, Thermo Fisher Scientific) and Alexa Fluor 568 goat anti-mouse IgG (1:500, Thermo Fisher Scientific) secondary antibodies in blocking buffer for 2 h at RT. Midguts were washed three times in 1xPBS, then mounted with ProLong®Diamond Antifade mountant with DAPI for visualization. To quantify TEP1 positive oocysts, 20 oocysts were randomly selected from individual mosquito midguts and the percentage displaying TEP1^+^ positive oocysts was recorded. Data were compiled from two independent experiments.

## Results

### Additional feeding differentially affects *Plasmodium* survival

To examine the effects of an additional blood meal on rodent malaria parasite infection, we first infected *An. gambiae* (Keele) with *Plasmodium berghei*, then maintained one cohort on sugar, while the second received a second, naive blood meal four days post-infection (Figure 1A). When oocyst numbers were examined at eight days post-infection, mosquitoes receiving a second blood meal displayed significantly reduced oocyst numbers when compared to those maintained on sugar alone after the initial infection (Figure 1B). To determine if this effect could be attributed to blood-feeding or the physical distention of the midgut that results from blood engorgement, we similarly challenged mosquitoes with *P. berghei* and maintained one cohort on sugar, while a second cohort was given a protein meal (2%BSA, 2mM ATP in 1xPBS) four days post-infection (Figure 1A). Following the additional protein feeding, oocyst numbers were similarly significantly reduced (Figure 1C). Based on the minimal components of a protein meal, we argue that the shared physical distention of the midgut of both treatments is responsible for this dramatic reduction in *P. berghei* numbers.

**Figure 1.**
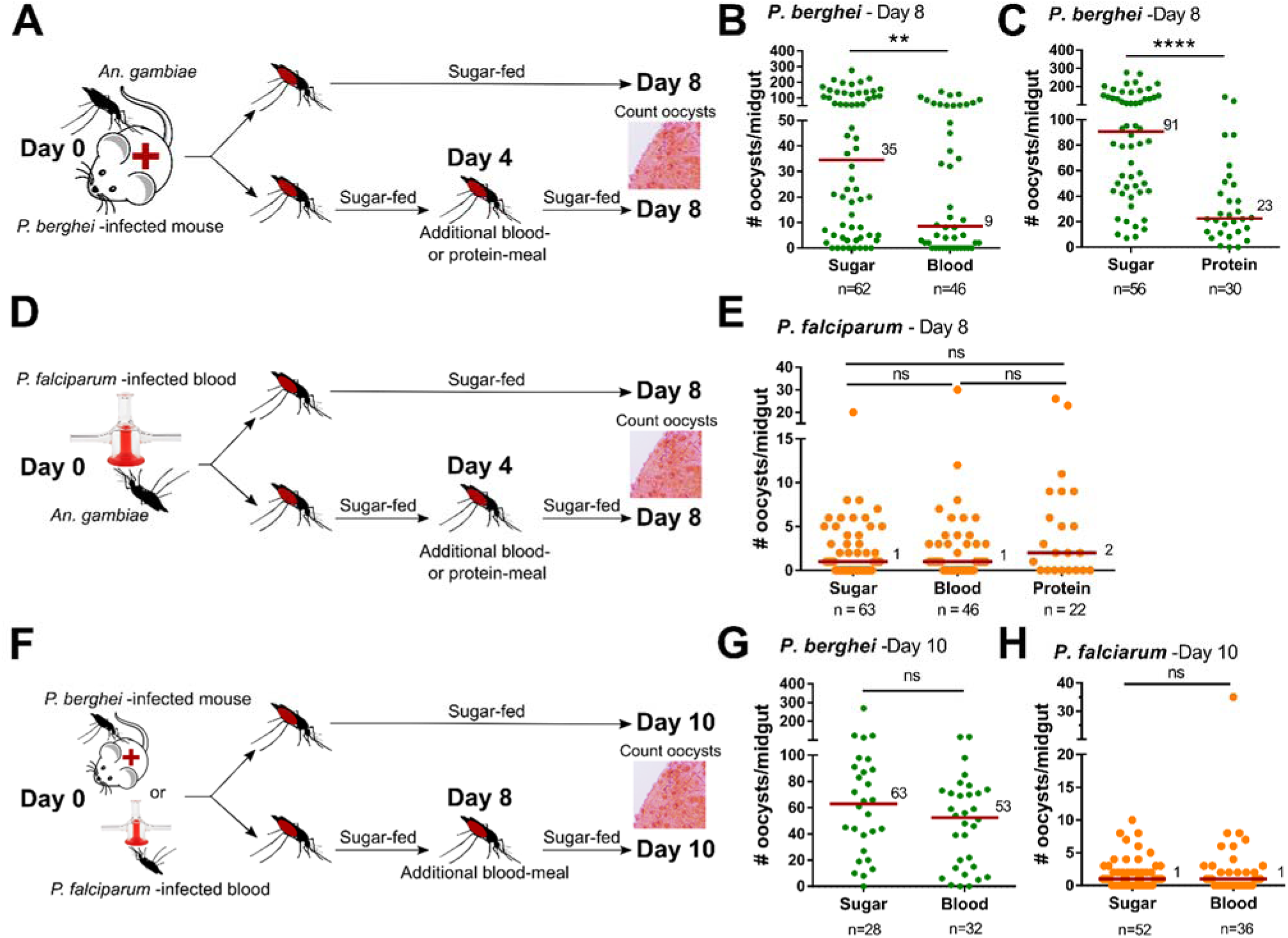
Additional feeding differentially impacts rodent and human malaria parasite survival. Experimental overview of *An. gambiae* feeding experiments where mosquitoes were initially challenged with *P. berghei*, then either maintained on sugar or received an additional uninfected blood- or protein-meal four days post infection (A). Oocyst numbers were examined at eight days post-infection for mosquitoes receiving an additional blood- (B) or protein-meal (C). Similar experiments were performed with *P. falciparum* (D), where oocyst numbers were evaluated at eight days post-infection following an additional blood- or protein-meal (E). To examine potential temporal effects on survival, experiments were outlined where mosquitoes were initially infected with *P. berghei* or *P. falciparum* and were maintained on sugar or received an additional uninfected blood-meal eight days post infection (F). Oocyst numbers were evaluated at ten days post-infection for *P. berghei* (G) and *P. falciparum* (H). For all experiments, each dot represents the number of parasites on an individual midgut, with the median value denoted by a horizontal red line. Data were pooled from 3 or more independent experiments with statistical analysis determined by Mann–Whitney analysis. Asterisks denote significance (\*\**P* < 0.01, \*\*\*\**P* < 0.0001). n = number of mosquitoes examined per group, ns = not significant.

Using a similar methodology, we also examined the influence of additional blood- or protein-meal on the infection of the human malaria parasite, *Plasmodium falciparum* (Figure 1D). However, an additional blood- or protein-meal did not influence *P. falciparum* oocyst numbers (Figure 1E), suggesting that there are differences in the immune recognition and killing of these two *Plasmodium* species in the mosquito host.

We also examined the temporal nature of an additional feeding, where mosquitoes infected with either *P. berghei* or *P. falciparum* received an additional blood-meal eight days post-infection (Figure 1F), a time in which developing oocysts have initiated sporogony (Smith and Barillas-Mury, 2016). At this stage of oocyst development, an additional blood meal does not influence *P. berghei* (Figure 1G) or *P. falciparum* (Figure 1H) oocyst numbers, which suggests there is a temporal component that determines *P. berghei* losses.

### Blood- and protein-feeding degrade the midgut basal lamina

*Plasmodium* oocysts develop in the space between the midgut epithelium and the midgut basal lamina (Smith and Barillas-Mury, 2016), providing protection from the cellular or humoral components of the mosquito immune system. To examine if an additional blood- or protein-meal could influence the integrity of the basal lamina, we utilized a collagen hybridizing peptide (CHP) that specifically binds unfolded collagen chains, to serve as an indicator of tissue damage (Hwang et al., 2017). Collagen IV serves as a primary component of the midgut basal lamina (Arrighi and Hurd, 2002; Arrighi et al., 2005; Dong et al., 2017) that becomes degraded following blood-feeding (Dong et al., 2017). We demonstrate that CHP stains dissected midguts shortly after blood-or protein-feeding (Figure 2A), with the intensity of CHP staining reaching peak levels ∼18hr after blood- (Figure 2B) or protein-feeding (Figure 2C), before being repaired shortly thereafter. Together, these results suggest the distention of the midgut and subsequent degradation of the basal lamina enable hemolymph immune components or host resources to interact with developing *Plasmodium* oocysts.

**Figure 2.**
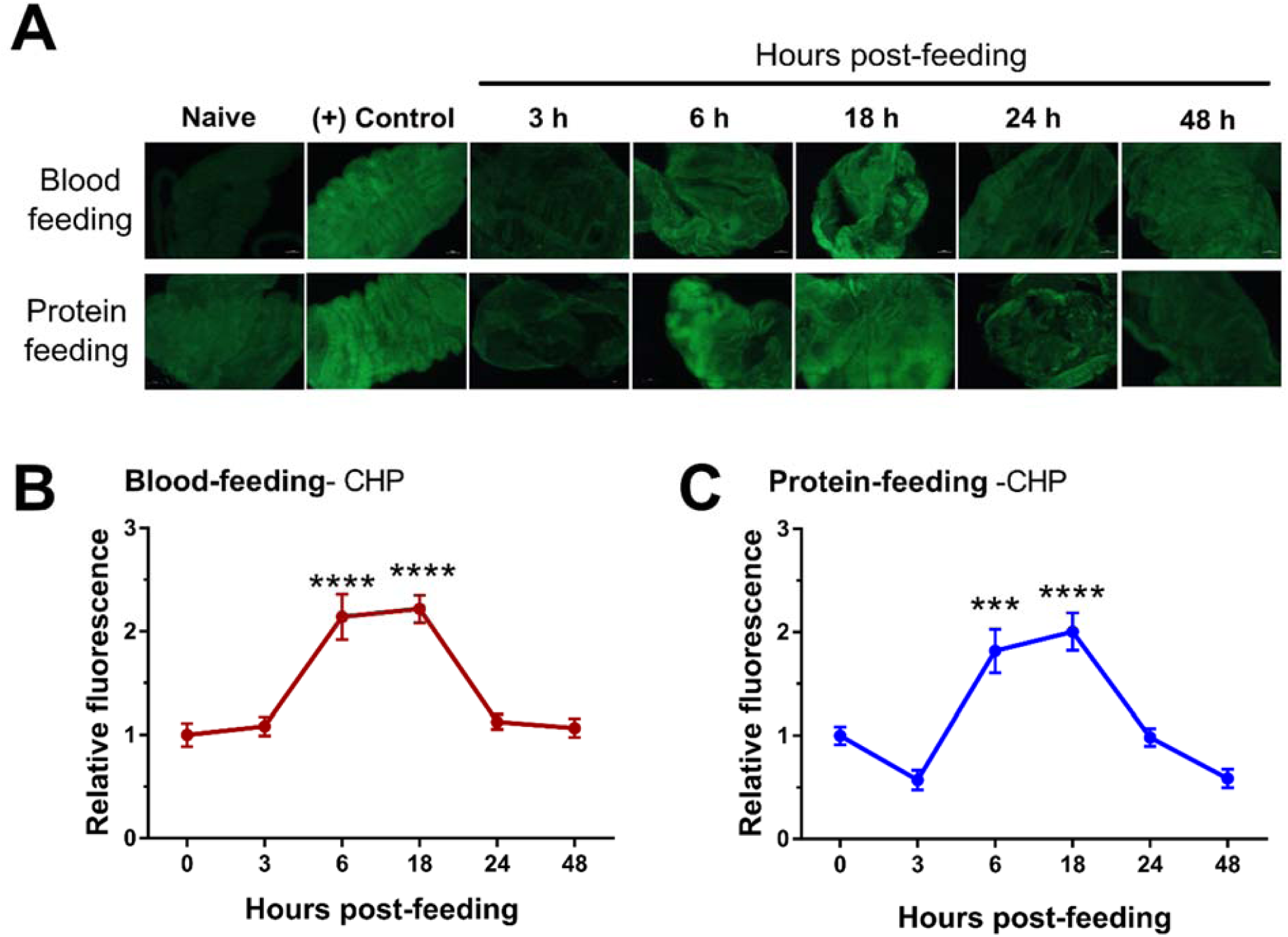
Mosquito feeding promotes the degradation of the midgut basal lamina. Using a fluorescein-labeled collagen hybridizing peptide (CHP) to detect degraded collagen, midgut basal lamina integrity was examined temporally at 3, 6, 18, 24 and 48 hours following blood- or protein-feeding (A). Heat-treated midguts (70° for 10 min. in 1x PBS) were used as a (+) control sample. The CHP fluorescence signal was quantified with Image J for each sample, and used to determine the relative fluorescence at each time point following blood-feeding (B) or protein-feeding (C). Significance was determined using a one-way ANOVA with a Holm-Sidak’s multiple comparisons test for comparison to the naïve (0 hr) timepoint. Asterisks denote significance (\*\*\**P* < 0.001, \*\*\*\**P* < 0.0001).

### Additional feeding enables TEP1-mediated killing of *P. berghei* oocysts

Since an additional feeding during the immature stages of *P. berghei* oocyst development limits survival (Figure 1) and feeding degrades the midgut basal lamina (Figure 2), we hypothesized that the decrease in *P. berghei* numbers could be attributed to the increased access of circulating immune components in the hemolymph to recognize developing parasites. To address this question, we examined the ability of TEP1, a major determinant of mosquito vector competence (Blandin et al., 2004; Fraiture et al., 2009; Povelones et al., 2009), to recognize the newly exposed *P. berghei* parasites following blood-feeding. Using immunofluorescence assays, we demonstrate that TEP1 recognition of *P. berghei* oocysts requires an additional blood-meal (Figure 3A), arguing that TEP1 may play an integral role in mediating these killing responses. We examined this further using a TEP1 knockout line of *An. gambiae* (Smidler et al., 2013), demonstrating that TEP1 is required for the losses in *P. berghei* oocyst numbers following an additional blood-feeding (Figure 3B). This argues that the degradation of the midgut basal lamina enables TEP1 to recognize and destroy *P. berghei* oocysts (Figure 3C), a stage of the parasite usually protected from mosquito complement recognition (Kwon et al., 2017; Smith et al., 2015).

**Figure 3.**
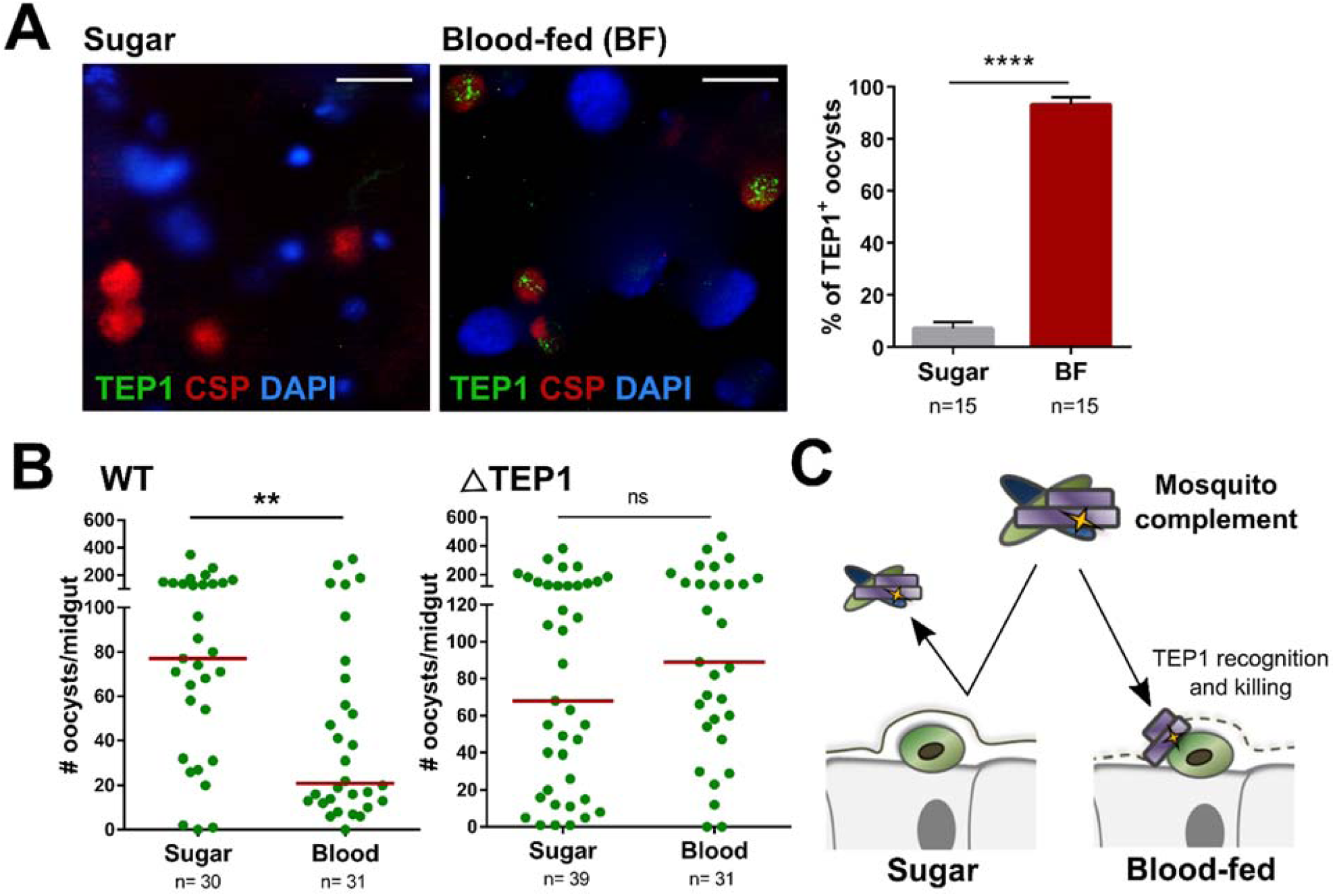
Additional feeding enables the recognition and killing of *P. berghei* oocysts by mosquito complement. Immunofluorescence assays were performed to examine TEP1 localization on developing oocysts when maintained on sugar or following an additional blood-meal. (A). Oocysts were identified by circumsporozoite protein (CSP) staining, enabling the ability to determine the percentage of TEP1^+^ oocysts from both experimental conditions. Additional feeding experiments were performed on either wild-type (WT) or mutant TEP1 (ΔTEP1) lines to confirm the involvement of mosquito complement in oocyst recognition and killing (B). Oocyst numbers were evaluated eight days post-infection. (C) Model for the role of mosquito complement via TEP1 recognition and killing of *P. berghei* oocysts. All results were examined by Mann-Whitney analysis where asterisks denote significance (\*\**P* < 0.01, \*\*\*\**P* < 0.0001). Scale bar: 10 μm. n = number of mosquitoes examined, ns = not significant.

### *P. falciparum* oocysts utilize host resources provided with an additional feeding

While an additional feeding does not limit human malaria parasite numbers unlike their rodent malaria counterparts (Figures 1 and 3), when evaluating oocyst numbers we noticed stark differences in parasite growth between *Plasmodium* species in the surviving oocysts (Figure 4). *P. falciparum* oocysts are significantly larger when mosquitoes receive an additional blood- or protein-meal when compared to mosquitoes maintained on sucrose alone after the infectious bloodmeal (Figure 4A), arguing that human malaria parasites are able to utilize host resources to accelerate their growth as previously suggested (Costa et al., 2018; Werling et al., 2019). Differences in oocyst size between blood- or protein-meal were not significant (Figure 4A). In contrast to these results with *P. falciparum*, similar experiments with *P. berghei* did not influence oocyst size (Figure 4B), suggesting that rodent malaria parasites are unable to utilize the extra mosquito host resources provided with an additional blood meal (summarized in Figure 4C). Taken together, our results support a model in which the human malaria parasite has evolved with its natural vector to evade immune recognition and to utilize host resources to increase the likelihood of its transmission.

**Figure 4.**
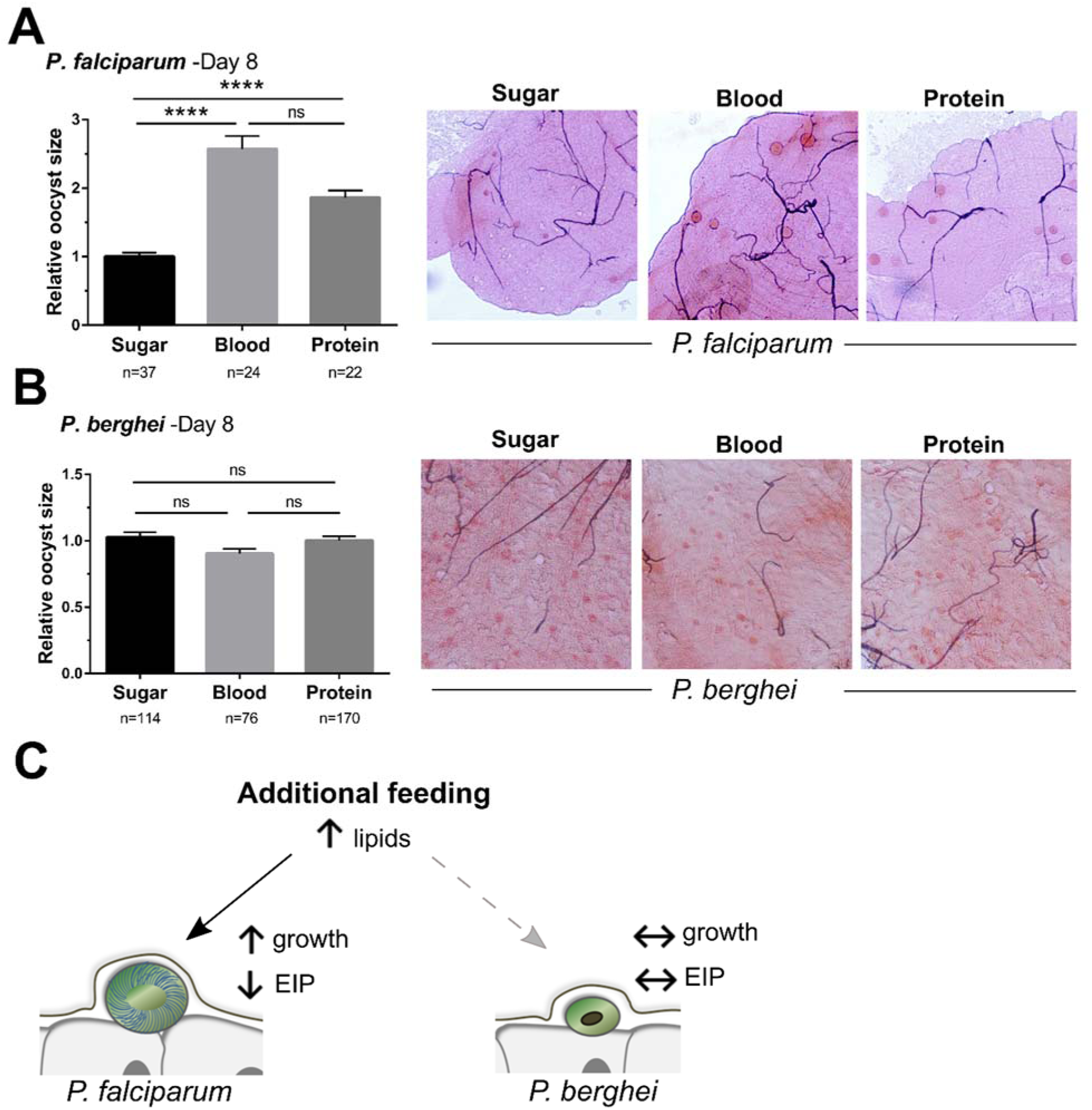
Human malaria parasites utilize host resources provided by an additional feeding to enhance their growth. *P. falciparum* (A) or *P. berghei* (B) oocysts were examined at eight days post-infection. The size of individual oocysts from mosquitoes maintained on sugar or that received an additional blood- or protein-meal were measured using Image J and compared by relative size across conditions. Representative images are shown on the right. Based on growth differences and supported literature, we propose a model in which human malaria parasites are able to utilize host resources to increase growth and increase the chances of transmission (C). Comparisons of oocyst size were analyzed using Kruskal-Wallis with a Dunn’s multiple comparison test. Asterisks denote significance (****P < 0.0001). n= number of oocysts examined, ns = not significant.

## Discussion

Studies of vectorial capacity in *Anopheles* have traditionally focused on measurements of *Plasmodium* oocyst or sporozoite numbers to evaluate the potential to transmit malaria. While insightful, these predominantly lab-based studies have not adequately addressed how mosquito physiology influences parasite survival and growth during the approximate 2-3 week extrinsic incubation period (EIP). Evidence suggests that larval nutrition (Shapiro et al., 2016) and temperature (Paaijmans et al., 2010; Shapiro et al., 2017) contribute to the EIP. Moreover, specific host-parasite interactions between different parasite and mosquito species have been suggested to influence vectorial capacity (Ohm et al., 2018; Simões et al., 2017), yet our current understanding of what defines these relationships remains limited. Since mosquitoes may feed multiple times during their lifespan, we hypothesized that an additional blood meal could influence parasite infection. Here we demonstrate that an additional blood meal significantly impacts *Plasmodium* development in terms of parasite survival and growth, with stark differences between human and rodent malaria parasites to evade immune recognition and to utilize nutrients provided by their mosquito host. These traits have likely evolved in *P. falciparum* through interactions with *An. gambiae* as its natural vector, while the laboratory model, *P. berghei*, has not been under similar selective pressures.

When challenged with an additional blood meal four days post-infection (the approximate time to potentially find a new host after completing a gonotrophic cycle), we see a dramatic reduction in the number of *P. berghei* oocysts, a phenotype recapitulated by similarly feeding on a protein meal. These results suggest that the effects of *P. berghei* killing are independent of the host blood meal, and more likely caused by the distention of the mosquito midgut following feeding. Ingestion of a blood- or protein-meal results in dramatic changes to midgut epithelium cell morphology, causing a flattening of the columnar cells, the loss of microvilli, and substantial degradation of the basal lamina (Dong et al., 2017; Sodja et al., 2007). This agrees with our CHP assays demonstrating the presence of degraded collagen shortly after taking a blood- or protein-meal, suggesting that the integrity of the basal lamina is compromised during distention, enabling the exposure of developing oocysts to components of the mosquito hemolymph.

Experiments with a TEP1 mutant line demonstrate the involvement of TEP1 and mosquito complement function in the killing of *P. berghei* oocysts following an additional blood-meal, contrasting previous results arguing that TEP1-mediated killing responses only target *Plasmodium* ookinetes (Blandin et al., 2004; Kwon et al., 2017; Smith et al., 2015). However, our data suggest that TEP1 can only recognize *P. berghei* early oocysts once the integrity of the basal lamina has been compromised by the distention of the midgut following an additional feeding. This supports a model in which the mosquito basal lamina serves as an integral physical barrier to protect the development of *P. berghei* from the mosquito innate immune system. In addition, these data argue that TEP1 recognition is temporally sensitive, where TEP1 binding and lysis is limited to the narrow time window before damage to the basal lamina is repaired at either the ookinete or oocyst stages of the parasite.

Therefore, it is of interest that an additional blood- or protein-meal does not similarly influence *P. falciparum* oocysts four days post-infection, suggesting that parasites surviving the transition into early oocysts can evade immune detection by mosquito complement. This is supported by previous work demonstrating the role of a parasite surface protein, P47, in complement evasion by *Plasmodium* ookinetes (Molina-Cruz et al., 2013, 2015; Ukegbu et al., 2017). As a result, we speculate that *P. falciparum* ookinetes able to evade complement recognition are also likely protected during the oocyst stage.

In addition to evaluating the effects of additional blood-feeding on parasite survival at four days post-infection, we also examined mature oocyst numbers when challenged at eight days post-infection once parasites have begun sporogony. However, at this later stage of parasite development, an additional blood-feeding did not affect either rodent or human malaria parasites. This suggests that there are differences in recognition of early and mature *P. berghei* oocysts, where only early oocysts are recognized by mosquito complement following an additional blood-meal. At present, why mature *Plasmodium* oocysts are no longer susceptible to killing remains unclear. This may be due to differences in immune detection, or alternatively, mature oocysts may have reached a stage of development in which they become immune privileged. There is support for the former due to the turnover of oocyst capsule proteins at the onset of sporogony (Smith and Barillas-Mury, 2016), which could potentially remove protein(s) involved in mosquito complement recognition of developing oocysts.

Based on differences in oocyst size observed during the initial evaluations of our infection experiments, we measured *Plasmodium* oocysts in response to an additional blood- or protein-meal. While no differences in *P. berghei* oocyst size were detected, *P. falciparum* oocysts significantly increased following additional feeding, suggesting that human malaria parasites utilize the added resources provided in a blood- or protein-meal to increase their growth. Previous work argues that the increase in lipid resources that accompany feeding are utilized by the developing oocyst for growth and sporozoite production (Costa et al., 2018; Werling et al., 2019). Moreover, an additional feeding increases the number of *P. falciparum* salivary gland sporozoites (Ponnudurai et al., 1989; Rosenberg and Rungsiwongse, 1991), arguing that this increased growth may enhance the potential for malaria transmission.

The impacts of an additional blood-meal were recently described in other vector-pathogen systems. In *Aedes aegypti* and *Ae. albopictus*, an additional blood-meal enhances arbovirus dissemination, increasing the transmission potential of ZIKV, DENV, and CHIKV (Armstrong et al., 2018). Moreover, sequential blood-feeding in sand flies leads to increased *Leishmania* parasite numbers and an improved frequency of transmission (Serafim et al., 2018). These examples suggest that blood-feeding is a conserved, yet relatively unexplored, mechanism for pathogens to decrease the EIP in their respective vector hosts.

Together, our experiments argue that human malaria parasites have developed the ability to evade immune detection and to utilize host resources in their natural mosquito vector. This in contrast to our experiments with rodent malaria parasites, which represent a non-natural system widely used in laboratory studies. This argues that *Plasmodium* species have evolved within their mosquito host, not only to evade immune detection as previously described (Collins et al., 1986; Molina-Cruz et al., 2012, 2015), but to also exploit resources provided with an additional blood-meal to accelerate their development and increase the chances of transmission. As a result, we believe our findings are an important advancement in our understanding of host-parasite interactions and the mechanisms that define vectorial capacity for the transmission of malaria.

## Acknowledgements

We would like to thank Doug Brackney for open discussions regarding this project and sharing protocols for CHP staining. We would also like to thank the Johns Hopkins Malaria Research Institute parasite and insectary core facility. This work was supported by the National Science Foundation Graduate Research Fellowship Program under Grant No. 1744592 to R.A.R, the Agricultural Experiment Station at Iowa State University to R.C.S, and by the National Institutes of Health, National Institute of Allergy and Infectious Diseases (R01AI122743 to G.D. and R21 AI44705 to R.C.S.).

## References

Armstrong, P.M., Ehrlich, H., Bransfield, A., Warren, J.L., Pitzer, V.E., and Brackney, D.E. (2018). Successive bloodmeals enhance virus dissemination within mosquitoes and increase transmission potential. BioRxiv 1–26.

Arrighi, R.B.G., and Hurd, H. (2002). The role of *Plasmodium berghei* ookinete proteins in binding to basal lamina components and transformation into oocysts. Int. J. Parasitol. 32, 91–98.

Arrighi, R.B.G., Lycett, G., Mahairaki, V., Siden-Kiamos, I., and Louis, C. (2005). Laminin and the malaria parasite’s journey through the mosquito midgut. J. Exp. Biol. 208, 2497–2502.

Blandin, S., Shiao, S.H., Moita, L.F., Janse, C.J., Waters, A.P., Kafatos, F.C., and Levashina, E.A. (2004). Complement-like protein TEP1 is a determinant of vectorial capacity in the malaria vector *Anopheles gambiae*. Cell 116, 661–670.

Collins, F.H., Sakai, R.K., Vernick, K.D., Paskewitz, S., Seeley, D.C., Miller, L.H., Collins, W.E., Campbell, C.C., and Gwadz, R.W. (1986). Genetic selection of a *Plasmodium*-refractory strain of the malaria vector *Anopheles gambiae*. Science 234, 607–610.

Costa, G., Gildenhard, M., Eldering, M., Lindquist, R.L., Hauser, A.E., Sauerwein, R., Goosmann, C., Brinkmann, V., Carrillo-Bustamante, P., and Levashina, E.A. (2018). Non-competitive resource exploitation within mosquito shapes within-host malaria infectivity and virulence. Nat. Commun. 9, 3474.

Dong, S., Balaraman, V., Kantor, A.M., Lin, J., Grant, D.A.G., Held, N.L., and Franz, A.W.E. (2017). Chikungunya virus dissemination from the midgut of *Aedes aegypti* is associated with temporal basal lamina degradation during bloodmeal digestion. PLoS Negl. Trop. Dis. 11, 1–26.

Ferguson, H.M., and Read, A.F. (2004). Mosquito appetite for blood is stimulated by *Plasmodium chaboudi* infections in themselves and their vertebrate hosts. Malar. J. 3, 12.

Fraiture, M., Baxter, R.H.G., Steinert, S., Chelliah, Y., Frolet, C., Quispe-Tintaya, W., Hoffmann, J. A., Blandin, S. A., and Levashina, E. A. (2009). Two mosquito LRR proteins function as complement control factors in the TEP1-mediated killing of *Plasmodium*. Cell Host Microbe 5, 273–284.

Gupta, L., Molina-Cruz, A., Kumar, S., Rodrigues, J., Dixit, R., Zamora, R.E., and Barillas-Mury, C. (2009). The STAT pathway mediates late-phase immunity against *Plasmodium* in the mosquito *Anopheles gambiae*. Cell Host Microbe 5, 498–507.

Hurd, H., Taylor, P.J., Adams, D., Underhill, A., and Eggleston, P. (2005). Evaluating the costs of mosquito resistance to malaria parasites. Evolution 59, 2560–2572.

Hwang, J., Huang, Y., Burwell, T.J., Peterson, N.C., Connor, J., Weiss, S.J., Yu, S.M., and Li, Y. (2017). *In situ* imaging of tissue remodeling with collagen hybridizing peptides. ACS Nano 11, 9825–9835.

Kwon, H., and Smith, R.C. (2019). Chemical depletion of phagocytic immune cells in *Anopheles gambiae* reveals dual roles of mosquito hemocytes in anti-*Plasmodium* immunity. Proc. Natl. Acad. Sci. 116, 201900147.

Kwon, H., Arends, B.R., and Smith, R.C. (2017). Late-phase immune responses limiting oocyst survival are independent of TEP1 function yet display strain specific differences in *Anopheles gambiae*. Parasit. Vectors 10, 369.

Molina-Cruz, A., DeJong, R.J., Ortega, C., Haile, A., Abban, E., Rodrigues, J., Jaramillo-Gutierrez, G., and Barillas-Mury, C. (2012). Some strains of *Plasmodium falciparum*, a human malaria parasite, evade the complement-like system of *Anopheles gambiae* mosquitoes. Proc. Natl. Acad. Sci. 109, E1957–E1962.

Molina-Cruz, A., Garver, L.S., Alabaster, A., Bangiolo, L., Haile, A., Winikor, J., Ortega, C., van Schaijk, B.C.L., Sauerwein, R.W., Taylor-Salmon, E., et al. (2013). The human malaria parasite Pfs47 gene mediates evasion of the mosquito immune system. Science 340, 984–987.

Molina-Cruz, A., Canepa, G.E., Kamath, N., Pavlovic, N. V, Mu, J., Ramphul, U.N., Ramirez, J.L., and Barillas-Mury, C. (2015). *Plasmodium* evasion of mosquito immunity and global malaria transmission: The lock-and-key theory. Proc. Natl. Acad. Sci. U. S. A. 112, 15178–15183.

Ohm, J.R., Baldini, F., Barreaux, P., Lefevre, T., Lynch, P.A., Suh, E., Whitehead, S.A., and Thomas, M.B. (2018). Rethinking the extrinsic incubation period of malaria parasites. Parasit Vectors 11, 1–9.

Paaijmans, K.P., Blanford, S., Bell, A.S., Blanford, J.I., Read, A.F., and Thomas, M.B. (2010). Influence of climate on malaria transmission depends on daily temperature variation. Proc. Natl. Acad. Sci. U. S. A. 107, 15135–15139.

Ponnudurai, T., Lensen, A.H.W., Van Gemert, G.J.A., Bensink, M.P.E., Bolmer, M., and Meuwissen, J.H.E.T. (1989). Sporozoite load of mosquitoes infected with *Plasmodium falciparum*. Trans. R. Soc. Trop. Med. Hyg. 83, 67–70.

Povelones, M., Waterhouse, R.M., Kafatos, F.C., and Christophides, G.K. (2009). Leucine-rich repeat protein complex activates mosquito complement in defense against *Plasmodium* parasites. Science 324, 258–261.

Ranford-Cartwright, L.C., McGeechan, S., Inch, D., Smart, G., Richterová, L., and Mwangi, J.M. (2016). Characterisation of species and diversity of *Anopheles gambiae* Keele colony. PLoS One 11, e0168999.

Rosenberg, R., and Rungsiwongse, J. (1991). The number of sporozoites produced by individual malaria oocysts. Am. J. Trop. Med. Hyg. 45, 574–577.

Schneider, C.A., Rasband, W.S., and Eliceiri, K.W. (2012). NIH Image to ImageJ: 25 years of image analysis. Nat. Methods 9, 671–675.

Serafim, T., Coutinho-Abreu, I. V, Oliviera, F., Meneses, C., Kamhawi, S., and Valenzuela, J.G. (2018). Sequential blood meals promote *Leishmania* replication and reverse metacyclogenesis augmenting vector infectivity. Nat. Microbiol. 3, 548–555.

Shapiro, L.L.M., Murdock, C.C., Jacobs, G.R., Thomas, R.J., and Thomas, M.B. (2016). Larval food quantity affects the capacity of adult mosquitoes to transmit human malaria. Proc. R. Soc. B Biol. Sci. 283.

Shapiro, L.L.M., Whitehead, S.A., and Thomas, M.B. (2017). Quantifying the effects of temperature on mosquito and parasite traits that determine the transmission potential of human malaria. PLoS Biol. 15, 1–21.

Simões, M.L., Mlambo, G., Tripathi, A., Dong, Y., and Dimopoulos, G. (2017). Immune regulation of *Plasmodium* is *Anopheles* species specific and infection intensity dependent. MBio 8, 1–13.

Smidler, A.L., Terenzi, O., Soichot, J., Levashina, E. A., and Marois, E. (2013). Targeted mutagenesis in the malaria mosquito using TALE nucleases. PLoS One 8, 1–9.

Smith, R.C., and Barillas-Mury, C. (2016). *Plasmodium* oocysts: overlooked targets of mosquito immunity. Trends Parasitol. 32, 979–990.

Smith, R.C., Vega-Rodríguez, J., and Jacobs-Lorena, M. (2014). The *Plasmodium* bottleneck: malaria parasite losses in the mosquito vector. Mem. Inst. Oswaldo Cruz 109, 644–661.

Smith, R.C., Barillas-Mury, C., and Jacobs-Lorena, M. (2015). Hemocyte differentiation mediates the mosquito late-phase immune response against *Plasmodium* in *Anopheles gambiae*. Proc. Natl. Acad. Sci. 112, E3412–20.

Sodja, A., Fujioka, H., Lemos, F.J.A., Donnelly-Doman, M., and Jacobs-Lorena, M. (2007). Induction of actin gene expression in the mosquito midgut by blood ingestion correlates with striking changes of cell shape. J. Insect Physiol. 53, 833–839.

Ukegbu, C.V., Giorgalli, M., Yassine, H., Ramirez, J.L., Taxiarchi, C., Barillas-Mury, C., Christophides, G.K., and Vlachou, D. (2017). *Plasmodium berghei* P47 is essential for ookinete protection from the *Anopheles gambiae* complement-like response. Sci. Rep. 7, 6026.

Werling, K., Shaw, W.R., Itoe, M.A., Westervelt, K.A., Marcenac, P., Paton, D.G., Peng, D., Singh, N., Smidler, A.L., South, A., et al. (2019). Steroid hormone function controls non-competitive *Plasmodium* development in *Anopheles*. Cell 177, 1–11.

WHO (2018). World Malaria Report. 2018. ISBN 978 92 4 156469 4.

